# Genome wide natural variation of H3K27me3 selectively marks genes predicted to be important for cell differentiation in *Phaeodactylum tricornutum*

**DOI:** 10.1101/2019.12.26.888800

**Authors:** Xue Zhao, Achal Rastogi, Anne Flore Deton Cabanillas, Ouardia Ait Mohamed, Catherine Cantrel, Berangère Lombard, Omer Murik, Auguste Genovesio, Chris Bowler, Daniel Bouyer, Damarys Loew, Xin Lin, Alaguraj Veluchamy, Fabio Rocha Jimenez Vieira, Leila Tirichine

## Abstract

- In multicellular organisms, Polycomb Repressive Complex2 (PRC2) is known to deposit H3K27me3 to establish and maintain gene silencing, critical for developmentally regulated processes. PRC2 complex is absent in both widely studied model yeasts which initially suggested that PRC2 arose with the emergence of multicellularity. However, its discovery in several unicellular species including microalgae questions its role in unicellular eukaryotes.
- Here, we use Phaeodactylum tricornutum enhancer of zeste E(z) knockouts and show that P. tricornutum E(z) is responsible for di and tri-methylation of lysine 27 of histone H3.
- H3K27me3 depletion abolishes cell morphology in P. tricornutum providing evidence for its role in cell differentiation. Genome wide profiling of H3K27me3 in fusiform and triradiate cells further revealed genes that may specify cell identity.
- These results suggest a role for PRC2 and its associated mark in cell differentiation in unicellular species and highlight their ancestral function in a broader evolutionary context than is currently appreciated.

## Introduction

Tri-methylation of lysine 27 of histone H3 (H3K27me3) is a mark deposited by Polycomb Repressive Complex 2 (PRC2), which mediates silencing of gene expression during differentiation and development in both animals and plants (Aldiri & Vetter, 2012, Fragola, Germain et al., 2013, Surface, Thornton et al., 2010). PRC2 is comprised of four core proteins, highly conserved among multicellular organisms: the histone methyltransferase (HMTase) enhancer of zeste *E(z)*, the WD40 domain containing polypeptide Extra Sex Comb *Esc*, the C2H2 type zinc finger protein Suppressor of zeste 12 *Su(z)12* and the Nucleosome remodeling factor 55 kDa subunit *Nurf-55* (Martinez-Balbas, Tsukiyama et al., 1998, Schwartz & Pirrotta, 2013). The absence of PRC2 in the unicellular yeast models *Saccharomyces cerevisiae* and *Schizosaccharomyces pombe* initially led to suggestions that it arose to regulate cell differentiation in multicellular organisms (Kohler & Villar, 2008). This hypothesis has recently been questioned because components of PRC2 and the associated mark H3K27me3 are found in several unicellular species that belong to different lineages, thus questioning the function of such a widespread complex in single celled organisms.

Only, few studies attempted to decipher the role of H3K27me3 in unicellular species. Using mass spectrometry, Shaver et al successfully identified mono- and di-methylation of lysine 27 on histone H3 in *Chlamydomonas reinhardtii* (Shaver, Casas-Mollano et al., 2010), however attempts to understand the role of H3K27me3 in this green alga were not conclusive because tri-methylation of lysine 27 could not be assessed reliably, due to either its absence or its nominal mass which could not be distinguished from acetylation of lysine 27 of the same histone(Shaver et al., 2010). In the model red alga *Cyanidioschizon merolae*, H3K27me3 was found to target genes and transposable elements. Genome-wide analysis revealed that H3K27me3 marked genes were related to organismal process, development, protein maturation and splicing (Mikulski, Komarynets et al., 2017). Recently, in another unicellular species, *Paramecium tetraurelia, E(z)-like* protein Ezl1 which contains the catalytic SET domain, was shown to be responsible of tri-methylation of both lysine 9 and 27 of histone H3 and that loss of both marks results in global derepression of transposable elements with modest effects on protein-coding gene expression (Frapporti, Miro Pina et al., 2019). These studies point to the importance of understanding H3K27me3 function in single celled species which was overlooked to decipher its role in an evolutionary context.

The pennate diatom *Phaeodactylum tricornutum (P. tricornutum*) which belongs to the stramenopile group of eukaryotes, only distantly related to the animal (Opisthokonta) and plant (Archaeplastida) eukaryotic crown groups has different morphotypes, fusiform (FM hereafter), which is the most prevailing morphology among the sampled accessions known so far, triradiate (TM hereafter), oval (OM hereafter) and cruciform (CM hereafter) (De Martino, Bartual et al., 2011, De Martino, 2007, He, Han et al., 2014, Rastogi, Vieira et al., 2020). Each morphotype can switch reversibly into a different morphology in response to several growth and environmental cues (De Martino et al., 2011). FM is the most stable morphotype while switching is more prominent in TM, CM and OM, which tend to convert to FM in the growth conditions used in this study (De Martino et al., 2011, De Martino, 2007).

Cell differentiation is often orchestrated by H3K27me3-mediated silencing that underlies the establishment and maintenance of cellular identity in multicellular model species (Margueron & Reinberg, 2011). Identification of H3K27me3 by mass spectrometry (Veluchamy, Rastogi et al., 2015) as well as the unique pleomorphic availability of *P. tricornutum* among stramenopiles present therefore an opportunity to understand its role in single celled organisms with respect to its potential contribution to establish morphotype switches as well as its function in an evolutionary context.

Our study points to the emerging function of PRC2 and its H3K27me3 associated mark as a determinant of the establishment and maintenance of cell morphology in the single celled species *P. tricornutum*, which shows signs of differentiation of the cell into diverse morphologies. This same function likely diversified with the emergence of multicellularity with PRC2 orchestrating development in plants and animals. This is the first evidence of the involvement of H3K27me3 in cell differentiation in a unicellular eukaryote only distantly related to animals and plants.

## Methods

### Strains and growth conditions

*Phaeodactylum tricornutum* Bohlin Clone Pt1 8.6 (CCMP2561) (referred as FM) and Clone Pt8Tc (referred as TM) cells were grown as described previously (Siaut, Heijde et al., 2007).

### Isolation and immunoprecipitation of chromatin

Chromatin isolation and immunoprecipitation were performed as described previously (Lin, Tirichine et al., 2012). The following antibodies were used for immunoprecipitation: H3K27me3 (07-449) from Millipore and H3K27me3 from cell signaling technology. qPCR on recovered DNA was performed as described previously (Lin et al., 2012)

### CRISPR/Cas9 plasmid construction

hCAS9n (Cas9 from *Streptococcus pyogenes*, adapted to human codon usage, fused to SV40 nuclear localization sequence, and containing a D10A mutation) was amplified from pcDNA3.3-TOPO-hCAS9n (kindly received from Dr. Yonatan B. Tzur, Hebrew University of Jerusalem), using the primers 5’-CAC CAT GGA CAA GAA GTA CTC-3’ and 5’-TCA CAC CTT CCT CTT CTT CTT-3’. The PCR product was first cloned into pENTR using pENTR/D-TOPO cloning kit (ThermoFisher Scientific), and then sub-cloned into a *P. tricornutum* pDest, containing an N-terminal HA-tag (Siaut et al., 2007), following the manufacturer’s protocol, which was named pDest-HA-hCAS9n.

For the sgRNA vector we first cloned the snRNA U6 promoter(Rogato, Richard et al., 2014) from *P. tricornutum* genomic DNA using the primers 5’-AAA CGA CGG CCA GTG AAT TCT CGT TTC TGC TGT CAT CAC C-3’ and 5’ - TCT TTA ATT TCA GAA AAT TCC GAC TTT GAA GGT GTT TTT TG-3’. PU6::unc-119_sgRNA (kindly received from Dr. Yonatan B. Tzur) backbone was amplified using the primers 5’-CAA AAA ACA CCT TCA AAG TCG GAA TTT TCT GAA ATT AAA GA-3’ and 5’-GGT GAT GAC AGC AGA AAC GAG AAT TCA CTG GCC GTC GTT T-3’. The two PCR products were used as template for a second round fusion PCR reaction as described in (Hobert, 2002). We further transformed the resulting product into *E. coli*, and extracted the ligated plasmid. The terminator sequence of the *P. tricornutum* U6 was amplified using the primers 5’-CATTCTAGAAGAACCGCTCACCCATGC-3’ and 5’-GTTAAGCTTGAAAAGTTCGTCGAGACCATG-3’, digested by XbaI/HindIII and ligated into XbaI/HindIII digested pU6::unc-119. The resulting vector, ptU6::unc-119-sgRNA, was used as template to replace the target sequence to *E(z)* target by PCR using primers 32817TS12fwd GTG TCG GAG CCC GCC ATA CCG TTT TAG AGC TAG AAA TAG C and 32817TS12rev GGT ATG GCG GGC TCC GAC ACC GAC TTT GAA GGT GTT TTT TG. Target sequences were picked using PhytoCRISP-Ex (Rastogi, Murik et al., 2016).

### Transformation of *P. tricornutum* cells and screening for mutants

Wild type cells of the reference strain FM and the TM were transformed with three plasmids (pPhat1, Cas9 and guide RNA with the target sequence) as described previously (Falciatore, Casotti et al., 1999). Positive transformants were validated by triple PCR screen for pPhaT1 shble primers (ACT GCG TGCACTTCGTGGC/TCGGTCAGTCCTGCTCCTC), sgRNA (GAGCTGGAAATTGGTTGTC/GACTCGGTGCCACTTTTTCAAGTT) and CAS9n (GGGAGCAGGCAGAAAACATT/TCACACCTTCCTCTTCTTCTT). For each colony, a rapid DNA preparation was performed as described previously and fragment of 400 bp was amplified with primers flanking the target sequence in the *E(z)* gene. The forward primer used is 5’-TAAGATGGAGTATGCCGAAATTC-3’ and reverse primer is 5’-AGGCATTTATTCGTGTCTGTTCG-3’ PCR product was run in 1% agarose gel and a single band was extracted using Machery Nagle kit and according to the manual manufacturer. PCR product was sequenced using the primer 5’-AGCCACCCTGCGTTAACTGAAAAT-3’.

To make sure that the fusiform cells originating from the switch of TM are not contaminants from the FM, each of the TM-T1, TM-Fusi and TM-N were checked for their genetic background whether it is FM or TM using a molecular marker designed around a 400 bp insertion in the FM background (Supplementary Fig. 3f), identified from genome sequencing of FM and TM strains of *P. tricornutum* (Rastogi et al., 2020). The PCR check confirmed that all the cell samples described above are in the TM genetic background.

### Validation of enrichment and expression of target genes

#### qPCR

Total RNA was extracted from TM and FM cells as described previously (Siaut et al., 2007) and cDNA was synthesized with cDNA high yield synthesis kit according to the manufacturer user manual. Quantitative PCR was performed as described previously (Siaut et al., 2007) using the primer list in Table S1. Briefly, cDNA was synthetized from 1 μg RNA using High-Capacity cDNA Reverse Transcription Kit (catalogue number 4368813) from Fischer scientific and according to manufacturer instructions. 1μl of cDNA was used in the QPCR reaction with the LightCycler^®^ 480 SYBR Green I Master (catalogue number 04707516001) from Roche and according to manufacturer instructions. Two reference genes were used, Tata Box binding Protein and Ribosomal Protein Small subunit 30S for normalization **(Siaut et al., 2007)**.

#### ChIP-qPCR

Chromatin Immunoprecipitation (ChIP) was done as described previously(Lin et al., 2012), primers were designed on random selected genes marked either by TM or FM and were listed in Table S1. Input DNA were diluted 10 times before qPCR. Quantitaibe PCR was either performed with Roche LightCycler 480 on 1ul of IP, diluted input and mock DNA respectively.

### Proteomics and PRM Measurements

Three independent histone purifications recovered from FM wild type cells as well as *E(z)* knockout mutant FM-Del6 in FM genetic background were simultaneously separated by SDS-PAGE and stained with colloidal blue (LabSafe Gel Blue GBiosciences). Three gel slices were excised for each purification and in-gel digested by using trypsin/LysC (Promega). Peptide extracted from each set were pooled and analyzed by nanoLC-MS/MS using an Ultimate 3000 system (Thermo Scientific) coupled to a TripleTOFTM 6600 mass spectrometer (AB Sciex). Peptides were first trapped onto a C18 column (75 μm inner diameter × 2 cm; nanoViper Acclaim PepMapTM 100, Thermo Scientific) with buffer A (2/98 MeCN/H2O in 0.1% formic acid) at a flow rate of 2.5 μL/min over 4 min. Separation was performed on a 50 cm x 75 μm C18 column (nanoViper C18, 3 μm, 100Å, Acclaim PepMapTM RSLC, Thermo Scientific) regulated to 50°C and with a linear gradient from 1% to 30% buffet B (100 MeCN in 0.085% formic acid) at a flow rate of 400 nL/min over 90 min. The mass spectrometer was operated in PRM top30 high sensitivity mode with 100 ms acquisition time for MS1 and MS2 scans respectively with included precursor mass list for 600 sec (see TableS1)

### PRM Data Analysis

The PRM data were analyzed using Skyline version 3.7.0.11317 MacCoss Lab Software, Seattle, WA; https://skyline.ms/project/home/software/Skyline/begin.view, fragment ions for each targeted mass were extracted and peak areas were integrated. The peptide areas were log2 transformed and the mean log2-area was normalized by the mean area of peptide STDLLIR using software R version 3.1.0. On each peptide a linear model was used to estimate the mean fold change between the conditions, its 97.5% confidence interval and the p-value of the two sided associated t-test. The p-values were adjusted with the Benjamini-Hochberg procedure(Benjamini, 2001).

The mass spectrometry proteomics data have been deposited to the ProteomeXchange Consortium via the PRIDE(Vizcaino, Cote et al., 2013) partner repository with the dataset identifier PXD012347.

#### Western blot analysis

Chromatin was extracted from wild type as well as mutants of both TM and FM cells and western blot performed as described previously (Lin et al., 2012).

### Sequencing and computational data analysis

#### ChIP-Seq

Chromatin Immunoprecipitation (ChIP) was done with monoclonal cell cultures grown using single triradiate cell from Pt8 population (referred as TM). CHIP-Seq was performed as described previously (Veluchamy et al., 2015). Two replicates were performed and showed a good Pearson correlation (Supplementary Fig. 4 and 5). Raw reads were filtered and low quality read-pairs were discarded using FASTQC with a read quality (Phred score) cutoff of 30. Using the genome assembly published in 2008 as reference (Pt1 8.6), we performed reference-assisted mapping of filtered reads using BOWTIE. We then performed the processing and filtering of the alignments using SAMTOOLS and BEDTOOLS. SICER (Zang, Schones et al., 2009) was then used to identify significant enriched H3K27me3 peaks by comparing it with the INPUT. Differential H3K27me3 peak enrichment analysis between FM and TM backgrounds was also done using SICER-df plugin. Peaks with Padj < 0.05 differential enrichment or depletion were considered significant. Functional inferences were obtained by overlapping the differentially enriched peaks over structural annotations from Phatr3 genome annotation (Rastogi, Maheswari et al., 2018).

#### Whole genome sequencing (WGS)

Whole genome sequencing was performed using DNA extracted from monoclonal cell cultures grown using single triradiate cell taken from TM and FM accession, respectively. At least 6 μg of genomic DNA from each accession was used to construct a sequencing library following the manufacturer’s instructions (Illumina Inc.). Paired-end sequencing libraries with a read size of 100 bp and an insert size of approximately 400 bp were sequenced on an Illumina HiSeq 2000 sequencer at Berry Genomics Company (China) and Fasteris for FM and TM, respectively. Low quality read-pairs were discarded using FASTQC with a read quality (Phred score) cutoff of 30. Using the genome assembly published previously (Bowler, Allen et al., 2008), we performed reference-assisted assembly of all the accessions. We used BOWTIE (-n 2 −X 400) for mapping the high quality NGS reads to the reference genome followed by the processing and filtering of the alignments using SAMTOOLS and BEDTOOLS. For estimating the genetic diversity between FM and TM genome, GATK (McKenna, Hanna et al., 2010) configured for diploid genomes, was used for variant calling, which included single nucleotide polymorphisms (SNPs), small insertions and deletions ranging between 1 and 300 base pairs (bp). The genotyping mode was kept default (genotyping mode = DISCOVERY), Emission confidence threshold (-stand_emit_conf) was kept 10 and calling confidence threshold (-stand_call_conf) was kept at 30. The minimum number of reads per base to be called as a high quality SNP was kept at 4 (i.e., read-depth >=4x). SNPEFF was used to annotate the functional nature of the polymorphisms.

#### RNA sequencing (RNA-Seq)

Total RNA was extracted from FM, TM, and FM-Del6 (FM E(z) knockout) cell lines. RNA expression and differential gene expression analysis was performed using Eoulsan version 1.2.2 with default parameters (Jourdren, Bernard et al., 2012). Genes having at least 2 folds expression change with P-value < 0.05 were considered as significant different expressed genes (DEGs). The workflow used for the RNA sequencing analysis can be found as Table S1.

#### GO enrichment analysis

DAMA and CLADE were used for Gene Ontology analysis. GO categories were grouped by 3 different levels of expression according to a simple density clustering algorithm (also confirmed by iterative k-means clustering).

#### R value analysis

An Information theoretic approach was used to measure the significant variation in quantities of transcripts in different experimental conditions (Zambelli, Mastropasqua et al., 2018). We derived entropy measure using FPKM values of 5 RNA-seq data (RPKM-Pt186, RPKM-Pt8TC, RPKM-PT186ko, RPKM-Pt8TCko, RPKM-Pt3Ov). Entropy levels of H3K27me3 marked genes were shown in boxplot. Two sample T-test with 95% confidence was performed on the entropy values and p-values were marked on the plot.

### Data Availability

All data are available through NCBI Sequence Read Archive with accession number PRJNA565539.

## Results

### CRISPR/cas9 knockout of *E(z)* and H3K27me3 depletion abolish cell morphology in *P. tricornutum*

PRC2 is a conserved complex with different subunits and orthologues in several multi and unicellular model species (Figure1A). In silico annotation of polycomb complex members in the Marine Microbial Eukaryotic Transcriptome Sequencing Project (MMESTP) shows the conservation and wide distribution of PRC2 complex in unicellular species suggesting the importance of this complex and its associated mark H3K27me3 in early diverging species (Supplementary Fig. 1A-C). In line with in silico identification of PRC2 complex subunits, western blot analysis using a monoclonal antibody raised against H3K27me3 in different representative of the Stramenopile, Alveolates and Rhizaria super lineage (SAR) identified the mark supporting further the importance of H3K27me3 in single celled species (Supplementary Fig. 1D). In the model diatom *P. tricornutum*, PRC2 complex is simpler than other organism: only one homolog was found in each subunit of PRC2 (Figure 1A), which makes *P. tricornutum* an ideal model organism for studying PRC2 and its associated mark H3K27me3 in unicellular species. Interestingly, western blot analysis of H3K27me3 in four different morphotypes of *P. tricornutum* (FM, TM, CM and OM cells) revealed a positive correlation between the complexity of the morphology (branching of the cell) and the amount of H3K27me3. Quantification of western blot shows that H3K27me3 level is higher in both CM and TM cells compared to FM and OM cells (Supplementary Fig. 1E). In fact, the relative expression of E(z) which is the catalytic enzyme that deposits H3K27me3 also presents the same tendency: TM shows higher expression of E(z) than FM and OM (Supplementary Fig. 1F).

**Figure 1.**
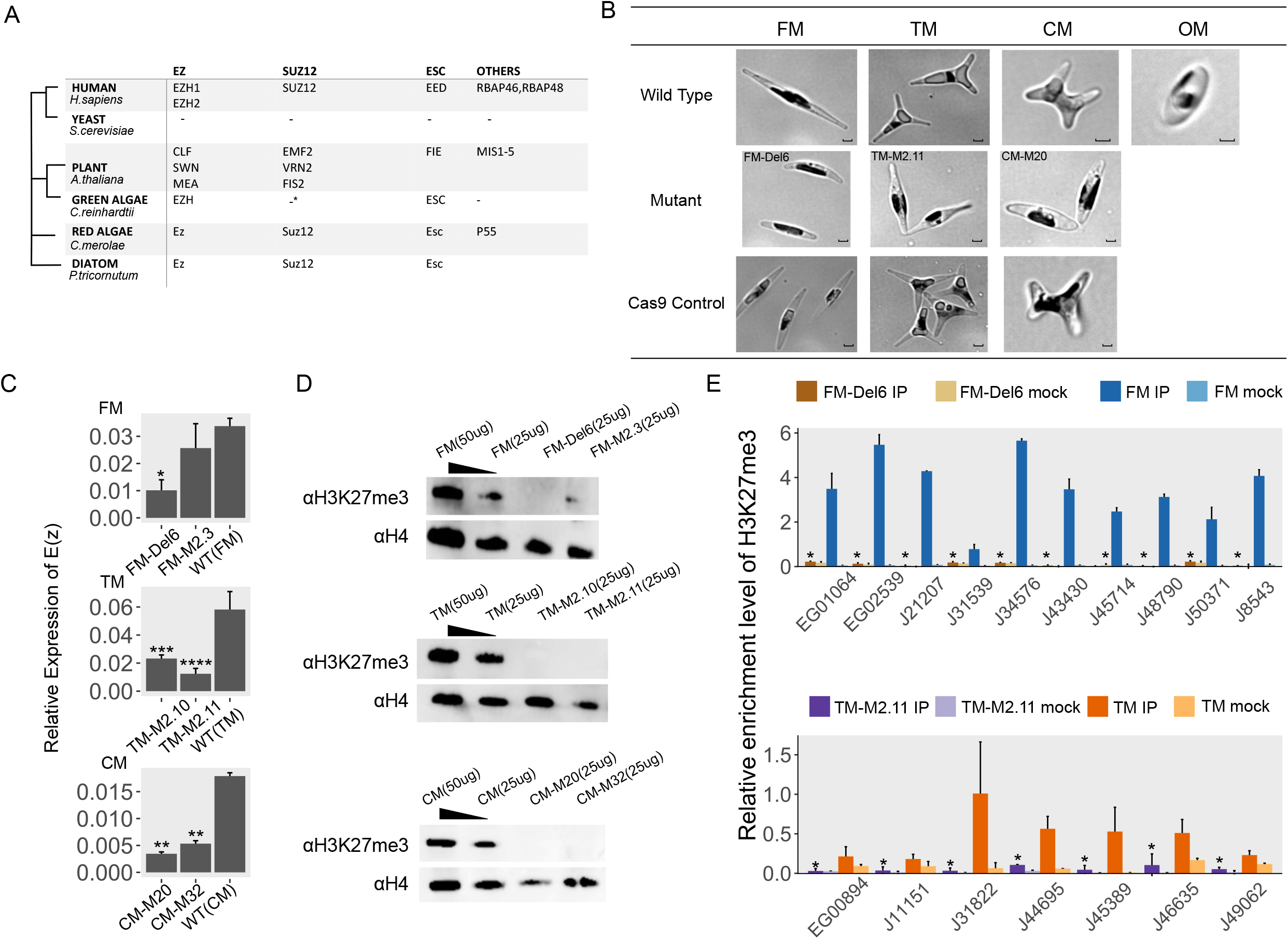
*Phaeodactylum tricornutum* morphotypes and enhancer of zeste knockout mutants. **(A)** Existence of Polycomb Repressive Complex2 (PRC2) complex components in model species including diatom. **(B)** Light microscopy images of wild type *Phaeodactylum tricornutum (P. tricornutum*) cells, *enhancer of zeste* E(z) Knockout(KO) lines and Cas9 control cells in four different kind of morphotype respectively. Scale bar of all WT, mutants and Cas9 control cells are 1 μm. Names of each mutant line were showed on the top left of their images.OM was the only one without E(z)KO line and Cas9 control due to lack of experimental capacities. **(C)** RNA expression levels of E(z) in wild type and E(z)KOs by qRT-PCR in three morphotypes background: fusiform (FM), triradiate (TM) and cruciform (CM) respectively (mean ± standard deviation; n = 4). Statistical analysis using Student t-test was performed by Mathematica. *P<0.05, **P<0.01, ***P<0.001, ****P<0.0001 **(D)** Western blots of Wild type (WT) and two E(z)KOs from each morphotype using a monoclonal antibody against H3K27me3. Histone H4 was used as a loading control. **(E)** ChIP-qPCR comparison of H3K27me3 enrichment levels (mean ± standard deviation; n = 4) on certain genes between WT and E(z) KO in FM and TM backgrounds. Statistical analysis using Student t-test was performed by Mathematica, average of P-value=0.000182672, *P<0.001.

To gain insights into the function of *E(z)* and its associated H3K27me3 mark in *P. tricornutum*, we generated two CRISPR/cas9 knockouts of *E(z)* in morphotypes (FM, TM and CM) leading to loss of function mutations including premature stop codon and frameshifts (Supplementary Fig1G). Light microscopy analysis of *E(z)* knockouts shows a change in cell morphology which becomes shorter in the FM background. Both triradiate and cruciform morphologies were abolished in TM and CM, respectively while transgenic lines carrying cas9 control vectors in each morphotype remain unchanged (Fig. 1B). Statistical analysis of cell size dimensions performed between FM, TM and their respective mutants show significant differences (Table S1). RNA expression of E(z) was significantly lower in E(z)KO lines except for one mutant FM-M2.3 which might due to incomplete removal of E(z) protein (Fig.1C). Knockout of *E(z)* also led to an overall depletion of H3K27me3 which we monitored by western blot using a monoclonal antibody against the mark (Fig. 1D), while *P. tricornutum* transgenic lines with the Cas9 control vector show similar H3K27me3 enrichment than the wild type (Supplementary Fig. 2A).

Mass spectrometry (MS) analysis of histones extracted from both wild type and *E(z)* knockout confirmed the loss of H3K27me3 and revealed a depletion of H3K27, demonstrating further the role of *E(z)* in tri-methylation of lysine 27 of histone H3 (Supplementary Fig.2B, C, D). Furthermore, MS analysis showed that di-methylation of lysine 27 is affected by the loss of *E(z)* (Table S1). Western blot assay of *E(z)* knockout mutants using a monoclonal antibody against H3K27me2 (Supplementary Fig.2E) confirmed the depletion of the mark from the mutants, supporting further MS study. Although cell shape is abolished in each of the mutants, knockout of *E(z)* is not a lethal mutation in *P. tricornutum*, only the growth of mutant lines are slightly slower compared to the wild type in each morphotype background. (Supplementary Fig.2F).

### Genome-wide distribution of H3K27me3 shows distinctive marked genes in fusiform and triradiate morphotype

To further investigate the role of H3K27me3 and its targets in different morphotypes, we carried out a genomic approach and performed Chromatin Immuno-Precipitation (ChIP) on two biological replicates of TM morphotypes using an antibody against H3K27me3 followed by DNA deep sequencing (ChIP-Seq) to generate a map of H3K27me3 distribution, which we compared to the one previously generated in FM in the same growth conditions (Veluchamy et al., 2015). ChIP-Seq data analysis revealed a similar H3K27me3 enrichment profile between TM and FM that localizes principally on transposable elements (TEs), with 58% and 60% of the reads overlapping with TE annotations for FM (Veluchamy et al., 2015) and TM, respectively(Supplementary Fig. 3A). The mark was found to occupy, on average, ~11.6% of the genome in FM cells, targeting approximately 15% of genes (consistent with (Veluchamy et al., 2015)) and ~13.2% of the genome within TM, targeting 19% of genes in agreement with the absolute amount of H3K27me3 to be elevated in TM compared to FM as detected by western blot (Supplementary Fig. 3B, Supplementary Fig. 1E). Indeed, more genes are marked by H3K27me3 in TM compared to FM, although most of the PRC2 targets are shared between both morphotypes (Fig. 2A) and exhibit globally broad coverage over the annotation (Supplementary Fig. 3C). TEs shared by FM and TM mainly belong to Copia type elements and include other categories such as Piggy Bac, SSR and MTITE type which seem to be equally distributed between the two morphotypes (Fig. 2B). Among the PRC2 targets, 635 and 297 genes are found to be specifically marked by H3K27me3 in TM and FM, respectively (Fig. 2A). We used ChIP followed by quantitative PCR (ChIP-qPCR) to validate H3K27me3 enrichment over specifically marked loci in both backgrounds, which corroborated the genome wide data for most of the tested genes (Fig. 2C). Commonly marked and unmarked loci were used as internal controls (Supplementary Fig. 3D). To test whether there are differences in nucleotide polymorphisms between marked, unmarked and total genes in both FM and TM that might contribute to morphotype switch, we compared single nucleotide polymorphism density between H3K27me3 marked and unmarked genes and found no significant differences (Supplementary Fig. 3E) supporting further a prominent role of H3K27me3 in regulating cell morphotypes in *P. tricornutum*.

**Figure 2.**
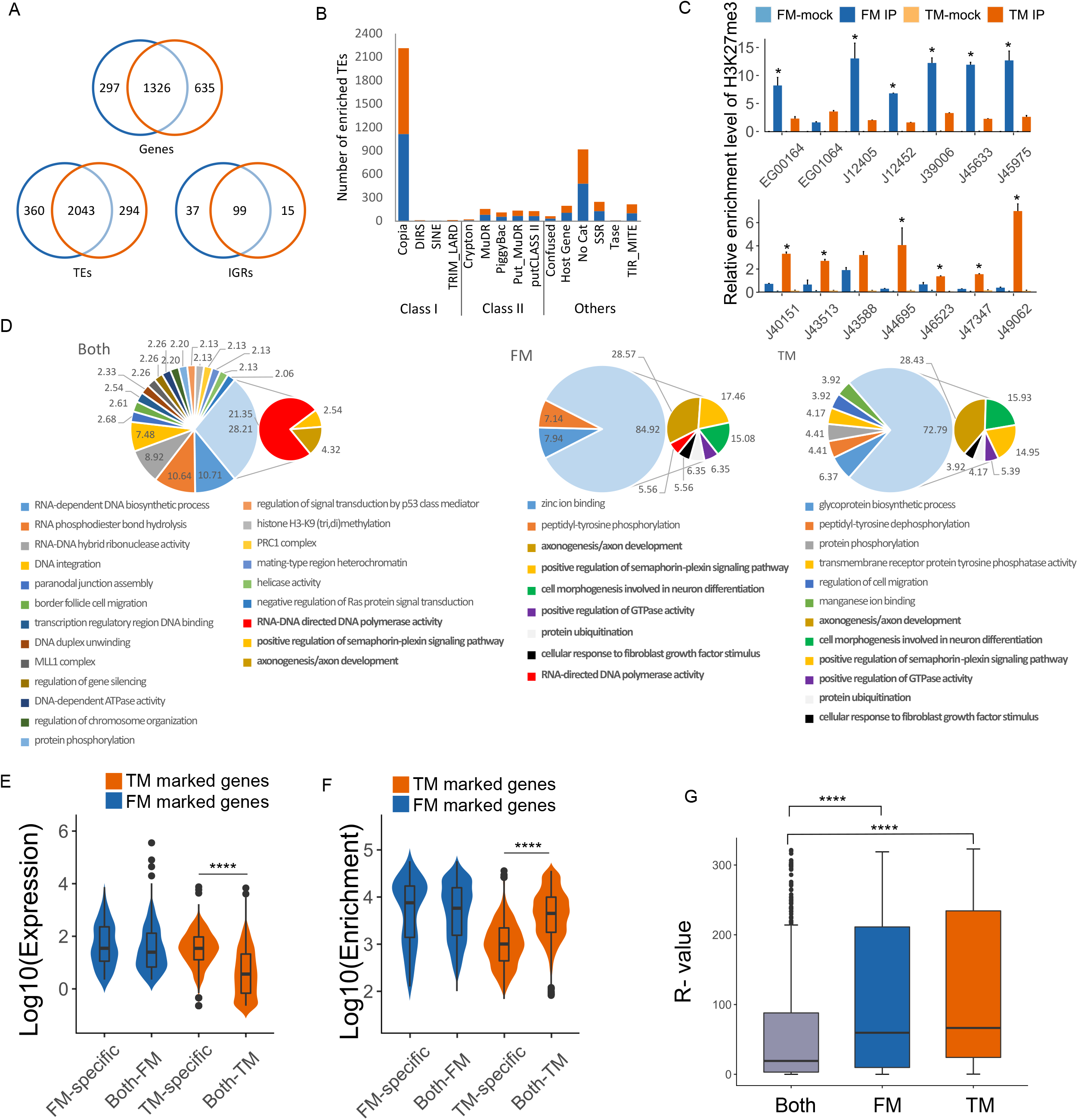
Genomic features of H3K27me3 targets in FM and TM cells in *Phaeodactylum tricornutum*. **(A)** Venn diagrams showing the number of common and specific genomic features [Genes, Transposable elements (TEs), and Intergenic Regions (IGRs)] targeted by H3K27me3 in TM (orange circles) and FM (blue circles). **(B)** Bar plot showing the distribution of TEs on different category in TM (orange bar) and FM (blue bar). **(C)** ChIP-qPCR validation of H3K27me3 specifically marked genes in TM and FM morphotypes (mean ± standard deviation; n = 4). Two experiment were conducted with different cell number so the enrichment level varies between two figures. **(D)** Distribution of the most frequent GO terms on genes marked with H3K27me3. The distribution was sub-divided into different categories, where TM and FM represent the GOs observed exclusively on triradiate and fusiform genes, respectively. A third category (Both) also presents a GO distribution for genes observed on both morphotypes. **(E)** Enrichment of H3K27me3 (Y-axis), with log10 scaling, over genes marked specifically in FM (blue), TM (orange), and also on genes marked in both (commonly marked) morphotypes. The enrichment profile is generated using number of genes marked by H3K27me3 specifically in each morphotypes and also in both. The significant H3K27me3 enrichment difference between specifically marked and commonly marked is estimated using two-tailed t-test with P value < 0.0001, denoted by “****”. **(F)** Expression of genes marked specifically in FM (blue), TM (orange), and in both phenotypic backgrounds with same principle aesthetics and categorical genes, used in 2E. The significance/non-significance of the variability of expression between specifically and commonly marked genes is estimated using two-tailed t-test with P value <0.0001, as denoted by “****”. “ns” denote non-significant. **(G)** Boxplot showing the entropy value distribution of H3K27me3 marked genes. Entropy values measures the differential expression of genes under different experimental conditions. Entropy values are derived from expression data (fragments per kilobase of exon per million fragments mapped) under five different experimental conditions. Genes marked specifically by H3K27me3 in TM (Triradiate) shows higher variation in expression followed by FM (Fusiform) specific H3K27me3 marked genes. Horizontal lines represent the median entropy values. P-values derived from t-test on FM/TM-specific H3K27me3 marked genes and genes marked on both conditions are shown on top.

To investigate the loss of H3K27me3 in *E(z)* knockout mutants, we performed ChIP-qPCR in FM and TM as well as the respective mutants and tested loci shown to be enriched by the ChIP-Seq analysis in FM or TM. We found a clear decrease in H3K27me3 enrichment over the tested loci in the mutants compared the FM and TM wild types which confirmed the depletion of H3K27me3 in these mutants as a consequence of *E(z)* loss (Fig. 1E).

### H3K27me3 specific targets in FM and TM are predicted to be important for cell differentiation

To gain insights into the functional categories enriched in H3K27me3 target genes that are shared between the two morphotypes or specific, we based our analysis on the updated Phatr3 annotation using DAMA (Bernardes, Vieira et al., 2016) and CLADE (Bernardes, Zaverucha et al., 2016) which is a machine learning methodology that uses pHMMs, position specific score matrix and supports vector machines to identify the corresponding most probable conserved regions to genes. DAMA and CLADE allow a more sensitive remote homology that permits to assign to conserved regions (even with no or poor conservation) the corresponding functional domains that would have been missed by other methods. From the functional domains identified, we obtained the corresponding GOs based on the method described in (Camon, Barrell et al., 2005), which has been evaluated at 91-100% accurate. To avoid obtaining general GOs, normally produced by other methods, we used eDAF (Zhao, 2020) to produce a more specific list of terms. Moreover, are considered statistically significant, only GO classes that are represented by at least 3 standard deviations above the average of observed entries.

Out of 1,640 H3K27me3 marked genes, 753 could not be assigned to a more specific GO category and are therefore marked as unknown. The genes that are marked in both morphotypes show top enrichment in RNA related biological processes such as RNA-dependent DNA biosynthetic process, RNA phosphodiester bond hydrolysis and RNA-DNA hybrid ribonuclease activity. Genes marked by H3K27me3 specifically in TM displayed top enrichment exclusively in (1) glycoprotein biosynthetic processes, (2) Peptidyl tyrosine dephosphorylation processes with Ankyrin repeats proteins (Fig. 2D, Table S1). Genes that are specifically marked in FM cells exhibit enrichment in exclusively categories such as peptidyl-tyrosine phosphorylation which is interestingly contrasting with the Peptidyl tyrosine dephosphorylation in TM; zinc ion binding proteins with a catalytic, co-catalytic or structural role known to be essential for the growth, development and differentiation of all living organisms(McCall, Huang et al., 2000) (Fig. 2D, Table S1). Interestingly, additional genes specifically marked in each FM or TM share categories with predicted functions in (1) positive regulation of GTPase activity (2) protein ubiquitination, both shown to play a role in differentiation of different cell types(Thompson, Loftus et al., 2008). Other shared functional categories point to GOs with genes involved in processes that occur while relatively unspecialized cells are acquiring the specialized features that permit a specific fate in multicellular species such as cell morphogenesis involved in neuron differentiation and cellular response to fibroblast growth factor stimulus GO categories.

### The role of H3K27me3 as a repressive mark is conserved

H3K27me3 is known to be a repressive mark in plants, animals and protists (Margueron & Reinberg, 2011, Shaver et al., 2010, Veluchamy et al., 2015). To further test the conserved role of H3K27me3 in repression in *P. tricornutum*, we analyzed the expression of genes in FM using whole genome transcript sequencing which revealed overall lower expression levels of H3K27me3-marked genes (Supplementary Fig 3F). Interestingly, when genes are marked by H3K27me3 in both FM and TM, their expression is lower compared to the genes that are uniquely marked in FM and to a lesser extent in TM (Fig. 2E). In line with result, enrichment level of H3K27me3 is significantly higher in common marked genes than specifically marked genes in TM (Fig. 2F).

We further analysed the effect of *E(z)* knockout on gene expression. Therefore, RNA sequencing of two biological replicates of the *E(z)* mutant (FM-Del6) was carried out and compared to previously generated RNA-seq in the wild type (FM). Around 1/4 of all genes are (23%, 2795 out of 12152 annotations) differentially expressed in the *E(z)* mutant (P-value. < 0.05), (Supplementary Fig. 3G, Table S1), indicating an essential role in gene regulation by PRC2 in *P. tricornutum*.

As a further test, we monitored by RT-qPCR the expression of randomly chosen 27 specifically marked genes in the *E(z)* knockout of the TM and found that 18 genes out of 27 showed a gain of expression in the mutant compared to the TM background, demonstrating further that depletion of H3K27me3 likely releases the repression of target genes and correlates with the loss of the triradiate morphology (Supplementary Fig.3H). There are 9 remaining genes including 2 with no change and 7 genes show more repression, suggesting that other gene expression regulation mechanisms might occur compensating for the loss of H3K27me3.

In order to find out whether the genes that are specific targets of H3K27me3 in either FM or TM show significant variations in expression across multiple growth conditions compared to marked genes in both morphotypes, we applied the R value analysis (Zambelli et al., 2018) which reflects the entropy, and therefore the variability in expression of genes between the above mentioned categories. The R value is used as an indication of genes differentially expressed in different conditions, the higher the value, the more differentially expressed is the gene. Our Analysis showed a higher R value in specifically marked genes compared to commonly marked ones (Fig. 2G).

### H3K27me3 dynamics correlates with morphotype switch

To substantiate the assumption that phenotypic plasticity and morphotype switch are regulated by PRC2 in *P. tricornutum*, we took advantage of the lack of stability of the TM phenotype and its tendency to switch to FM. Specifically, we used clonal cell samples with FM and TM morphologies from the same genetic background (TM), which switches habitually to fusiform and therefore contains a mixture of FM and TM cells. Molecular markers were chosen to distinguish TM and FM strains and exclude any cross contamination (Supplementary Fig.3I). We reasoned that the activity of *E(z)* should correlate with H3K27me3 levels in the following way: (1) a pure triradiate population isolated from TM-N (named here TM-T1): highest level of H3K27me3), (2) a population of cells from TM after N generations (N is 60 ± 5) of cell division containing a mixture of triradiate and fusiform morphotypes (TM-N): medium level of H3K27me3) and (3) fusiform cells isolated from the triradiate background TM (TM-Fusi): lowest level of H3K27me3 (Fig. 3A). We performed a quantitative PCR analysis of *E(z)* transcript levels which shows a clear decrease in TM-N and TM-Fusi compared to TM-T1 (Fig. 3B). These results correlate with the switch from TM to FM, reflecting a lower activity of E(z) and H3K27me3 levels.

**Figure 3.**
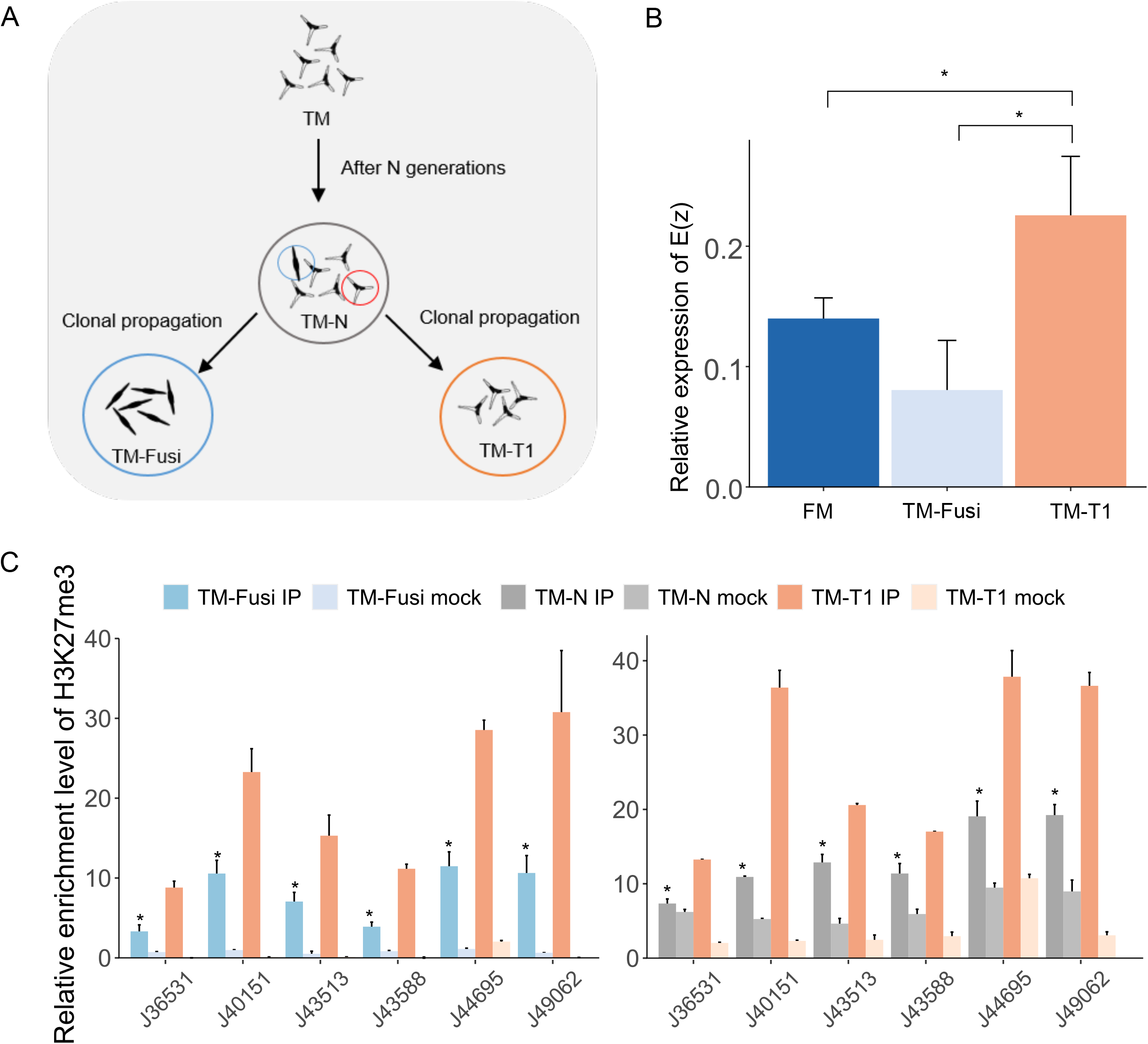
H3K27me3 enrichment levels and morphotype switch in *Phaeodactylum tricornutum*. **(A)** Schematic diagram showing generation of cell lines used in morphotypr switch experiment. After generations of culture in lab growth condition (ASW media, 19°C, 12h/12h light dark period), some triradiate cells switch to fusiform forming a mixture of FM and TM cells, named **TM-N**. A single fusiform cell from TM-N was picked and propagated clonally giving rise to **TM-Fusi**. Similarly, single triradiate cell was isolated from TM-N and its clonal propagation gave a population of pure triradiate cells named **TM-T1**. (**B)** Relative expression level of Enhancer of Zest in FM, TM-T1 and TM-Fusi respectively (mean ± standard deviation; n = 4) (**C)** ChIP-qPCR shows enrichment levels of H3K27me3 between TM-T1 and TM-Fusi, and also betweenTM-T1 and TM-N on certain H3K27me3 marked loci (mean ± standard deviation; n = 4).

We then asked whether specifically H3K27me3-marked loci in TM lose the mark upon cell switching to FM after multiple generations of sub-culturing leading to TM-N. Likewise, we asked whether fusiform cells (TM-Fusi) resulting from the switching from TM contain the same level of H3K27me3 enrichment than FM. As expected, ChIP-qPCR analysis showed clearly a loss of the mark in a population containing a mixture of fusiform and triradiate cells (TM-N) as well as in TM-Fusi compared to TM-T1 (Fig. 3C, D), which contains only triradiate cells, thus correlating the morphology with the level of enrichment in H3K27me3 over specific genes.

## Discussion

Polycomb proteins originally identified in *Drosophila* were shown to be important for the regulation of the spatiotemporal expression of transcription factors, most notably the Hox genes, themselves important for body segmentation in the fly. Since then, many orthologous were identified in plants and animals with similar functions, namely the orchestration of proper cell fate identity and development (Shaver et al., 2010). PRC2 is the best characterized complex with the enhancer of zeste catalytic subunit which is involved in tri-methylation of lysine 27 of histone H3. Knockout of this enzyme leads to cell fate modifications and developmental defects in plants and animals(Margueron & Reinberg, 2011). Although, widely studied in multicellular species, PRC2 complex and its associated mark H3K27me3 are scarcely identified in unicellular organisms and their role poorly investigated in particular in cell fate determination where to our best of knowledge, no evidence for such a role was reported so far in single celled species. The absence of the complex in the most studied yeast models species, *Saccharomyces cerevisiae* and *Schizosaccharomyces pombe* led to the assumption that PcG complexes arose with the emergence of multicellularity. The availability of genome sequences(Keeling, Burki et al., 2014) of many single celled organisms belonging to different super group such as Chromalveolata, Opisthokonta and Archaeplastida (Supplementary Fig. 1A, B) revealed the existence of several components of the polycomb complex which supports the idea of its emergence in an ancestral unicellular eukaryote and its probable loss in specific lineages during evolution (Shaver et al., 2010). This raises the question of their role in single celled species knowing that in multicellular organisms PcG are involved in developmental processes that are not necessarily relevant to single celled species.

In the model diatom *P. tricornutum*, we have previously identified without ambiguity, H3K27me3 by mass spectrometry and investigated its distribution genome wide showing that it represses mainly transposable elements and a significant number of genes (Veluchamy et al., 2015). *P. tricornutum* can be found in four morphotypes, fusiform (FM), triradiate (TM), oval (OM) and cruciform (CM), all capable of switching into a different morphology reversibly and in a relatively short time suggesting chromatin mediated regulation of cell morphology. Interestingly, western blot analysis using a monoclonal antibody against the mark in those four morphotypes revealed a higher level of H3K27me3 in TM and CM which cells are more branched compared to FM and OM who have simpler morphology (Supplementary Fig. 1E), suggesting a positive correlation between the complexity of the morphology and the absolute quantity of H3K27me3, indicating a morphology dependent PRC2 activity in *P. tricornutum*. Likewise, expression of E(z) is significantly higher in TM compared to other morphotypes (Supplementary Fig. 1F), which further supports the hypothesis of H3K27me3 activity dependent morphology. *P. tricornutum*, with its conserved single copies of PRC2 complex subunits, the reliably identified H3K27me3 and the presence of different morphologies which allows phenotypic screen is a unique opportunity to investigate the role of PRC2 complex and its associated mark, H3K27me3 in unicellular species.

We used CRISPR cas9 editing to generate mutants in the homologue of the catalytic subunit of PRC2, *enhancer of zeste* E(z) in each of FM, TM and CM backgrounds leading to a total depletion of H3K27me3 shown by western blot analysis and mass spectrometry. This demonstrates the role of *P. tricornutum* E(z) as a histone methyltransferase responsible for tri-methylation of lysine 27 of histone H3. H3K27me3 depletion causing changes in the cell morphology of all the morphotypes in particular in TM and CM where the branching of the cell is completely abolished suggests that the loss or diminution of *E(z)* activity, and hence H3K27me3, induces the observed changes in cell morphology (Fig. 1B and D). More importantly, mutants of E(z) show a very stable morphotype, no changes were observed under culture conditions used in this study. This evidenced that *E(z)* and its associated mark are required for morphotype switch to establish specific cell identity in a unicellular species. Similar to animals, plants and *Neurospora crassa*, H3K27me3 appears not to be essential for cell survival in *P. tricornutum*, as indicated by the overall growth of E(z) knockout lines which are not severely affected, only slightly retarded compared to wild type lines (Supplementary Fig. 2F). Although E(z) knockout has in general shorter size than the wild type(Figure 1B, Table S1), the morphotype remains fusiform which is not similar to the stress-induced natural OM morphotype reported before (Ovide, Kiefer-Meyer et al., 2018).

Similar to fungi (Jamieson, Rountree et al., 2013) and mammals (Ferrari, Scelfo et al., 2014), *E(z)* was found to dimethylate lysine 27 of histone H3 in *P. tricornutum*. However, this is different from *A. thaliana* where PRC2 loss of function leads to specific depletion of H3K27me3(Lafos, Kroll et al., 2011), although in vitro assays with reconstituted *A. thaliana* PRC2 components showed mono, di and tri-methylation of lysine 27 of histone H3(Jacob, Feng et al., 2009, Schmitges, Prusty et al., 2011).

The pattern of distribution of H3K27me3 genome wide in TM is similar to FM targeting mainly transposable elements and more genes in TM than FM with a broad pattern of distribution as reported in our previous study (Veluchamy et al., 2015). In the unicellular species, *Cyanidioschizon merolae*, *Chlamydomonas reinhardtii* and *Paramecium tetraurelia*, H3K27me3 was also found to target primarily TEs suggesting that one of the ancestral roles of PCR2 may have been in defense responses against intragenomic parasites such as TEs, prior to being co-opted for lineage specific functions like developmental regulation in multicellular eukaryotes (Frapporti et al., 2019, Shaver et al., 2010). Although, most of H3K27me3 target genes are shared between FM and TM, some were found to be marked exclusively in either of the morphotypes suggesting a specific role of these genes in cell differentiation leading to a triradiate or a fusiform morphology.

Functional annotation of H3K27me3 target genes using a highly sensitive annotation tool, DAMA and CLADE allowed the assignation of GO categories which clearly distinguished between those found in commonly marked genes(noted as Both in Fig. 2D) from the GOs exclusively found in FM or TM (noted as FM and TM respectively in Fig. 2D). Genes which are marked commonly mainly belong to GO categories related to RNA/transcription activity, representative categories include: RNA-DNA directed DNA polymerase activity (21.35%), RNA-dependent DNA biosynthetic process (10.71%), RNA-DNA hybrid ribonuclease activity (8.92%) and DNA integration (7.48%). These categories together representing 48.46% of annotated GOs terms, reflect the importance of the mark throughout the cell cycle progression to ensure repression of specific genes and the transmission of the mark to the replicating fork which is in line with the function of H3K27me3 in other organism. Interestingly, presence of certain categories such as MLL1 complex (catalyzing methylation of H3K4), PRC1 complex (catalyzing methylation of H2AK119Ubi) and histone of H3-K9(di, tri) methylation in commonly marked genes suggests the possible correlation of H3K27me3 and other histone marks such as H3K4me3, H2AK119Ubi and H3K9me2/3. This finding is in line with the co-occurrence of these histone modifications previously reported in *P. tricornutum* (Veluchamy et al., 2015).

Interestingly, additional genes specifically marked in each FM or TM share categories with predicted functions in (1) Axonogenesis/axon development (28.57%), (2) positive regulation of semaphoring-plexin signaling pathway (17.46%), (3) cell morphogenesis involved in neuron differentiation (15.08%), (4) cellular response to fibroblast growth factor stimulus (5.56%), which all together represent 66.67% of the GOs suggesting the correlation of these specific marked genes with morphogenesis. Although these categories are known to be involved in general processes, they were reported in different studies with a role in morphogenesis. This is exemplified by : semaphorin-plexin signaling pathway related genes known to be directly related to cytoskeletal and adhesive machinery that regulate cellular morphology (Alto & Terman, 2017). GTPase positive regulation activity reported with a role in cell morphology changes, and neurite outgrowth and guidance (Etienne-Manneville & Hall, 2002) as well as the differentiation of many cell types, including neurons, T lymphocytes and myocytes (Bryan, Li et al., 2005); protein ubiquitination shown to play a role in the complex regulation of the levels and function of many proteins and signaling pathways involved in determining cell fate(Thompson et al., 2008)(Fig. 2D, Supplementary Table S1). Typical examples in TM specific associated GOs also include (1) glycoprotein biosynthetic processes involved in the transfer of sugar moieties that might determine different sugar composition of the cell wall which is known to be sugar rich in *P. tricornutum(Le Costaouëc, 2017*), (2) Peptidyl tyrosine dephosphorylation processes with Ankyrin repeats proteins known to act as scaffold for connecting molecular interactions, likely important for development of the numerous signaling pathways associated generally to more complex multicellular organisms (Marcotte, Pellegrini et al., 1999) (Fig. 2D, Supplementary Table S1). Genes that are specifically marked in FM cells exhibit enrichment in categories such as peptidyl-tyrosine phosphorylation containing genes with central roles as modulators of cell differentiation and cell fate decisions (Yu & Cui, 2016). Overall, the genes that are specifically marked in the TM or FM morphotypes reflect processes related to cell growth, proliferation and differentiation. Although some of the categories seem not relevant to single celled organisms, our GO analysis may reflect evolutionary vestige tracing back to a shared ancestor (Weiss, Preiner et al., 2018).

Gene expression levels which were found higher in H3K27me3 specifically marked genes compared to common targets (Fig 2E) suggest that exclusively marked genes in either FM or TM are kept under less stringent and tight repression which might be due to their putative role in morphotype switch, which is known to be a dynamic process (De Martino et al., 2011). R value analysis (Maheswari, Jabbari et al., 2010) which reflects the expression variation of these genes found to be higher in FM and TM specifically marked genes compared to shared H3K27me3 targets, which means that specifically marked genes are more actively expressed compared to commonly marked ones (Fig. 2G). It supports further the finding of exclusive H3K27me3 enrichment and higher expression versus shared enrichment and lower expression (Fig. 2E and F). RT-qPCR in TM-M2.11 (the knockout of *E(z)* in TM) versus TM suggests further that depletion of H3K27me3 likely releases the repression of target genes and correlates with the loss of the triradiate morphology (Supplementary Fig3 G). Although the remaining genes showed no change or even a gain in expression, these genes can be targets of other repressive or active marks.

The correlation between H3K27me3 enrichment and cell morphology is further substantiated with the dynamic of both *E(z)* expression and the mark enrichment when cells switch from TM to FM. Morphotype switch experiment clearly shows a higher expression of *E(z)* in pure triradiate compared to a mixture of FM and TM and pure FM cells, all three originating from the same TM genetic background (Fig.3B and C). H3K27me3 dynamic and changes in the morphology in an identical genetic background is a strong evidence for chromatin mediated regulation of cell differentiation in *P. tricornutum*.

In summary, we have demonstrated in this study the role in *P. tricornutum E(z*) as a histone methyltransferase responsible for di and tri-methylation of lysine 27 of histone H3. Knockout of *E(z)* causes H3K27me3 depletion and loss of triradiate cell shape maintenance, providing evidence for the involvement of *E(z)* and its associated mark in establishing and/or maintaining cell morphology in unicellular species. We showed the dynamic nature of the mark, depending on the specific morphology between and within *P. tricornutum* accessions that correlates with the level of H3K27me3 enrichment. We showed differential marking in two different accessions of *P*. *tricornutum*, FM versus TM, which identified genes related to cell fate decisions compared to commonly marked genes. This is the first evidence for the involvement of H3K27me3 in cell differentiation in a unicellular eukaryote only distantly related to animals and plants. Our study points to the emerging function of PRC2 and its H3K27me3 associated mark as a determinant of the establishment and maintenance of cell morphology in a single celled species such as *P. tricornutum* that shows signs of differentiation of the cell into diverse morphologies. This same function likely diversified with the emergence of multicellularity with PRC2 orchestrating development in plants and animals.

## Supporting Information

**Supplementary Table 1.** List of genes marked by H3K27me3 with their annotation and GOs. This table also includes in different sheets: the cell of different morphotype counts and cell size measurement, primer list and mass spectrometry quantification.

**Supplementary Figure 1.**
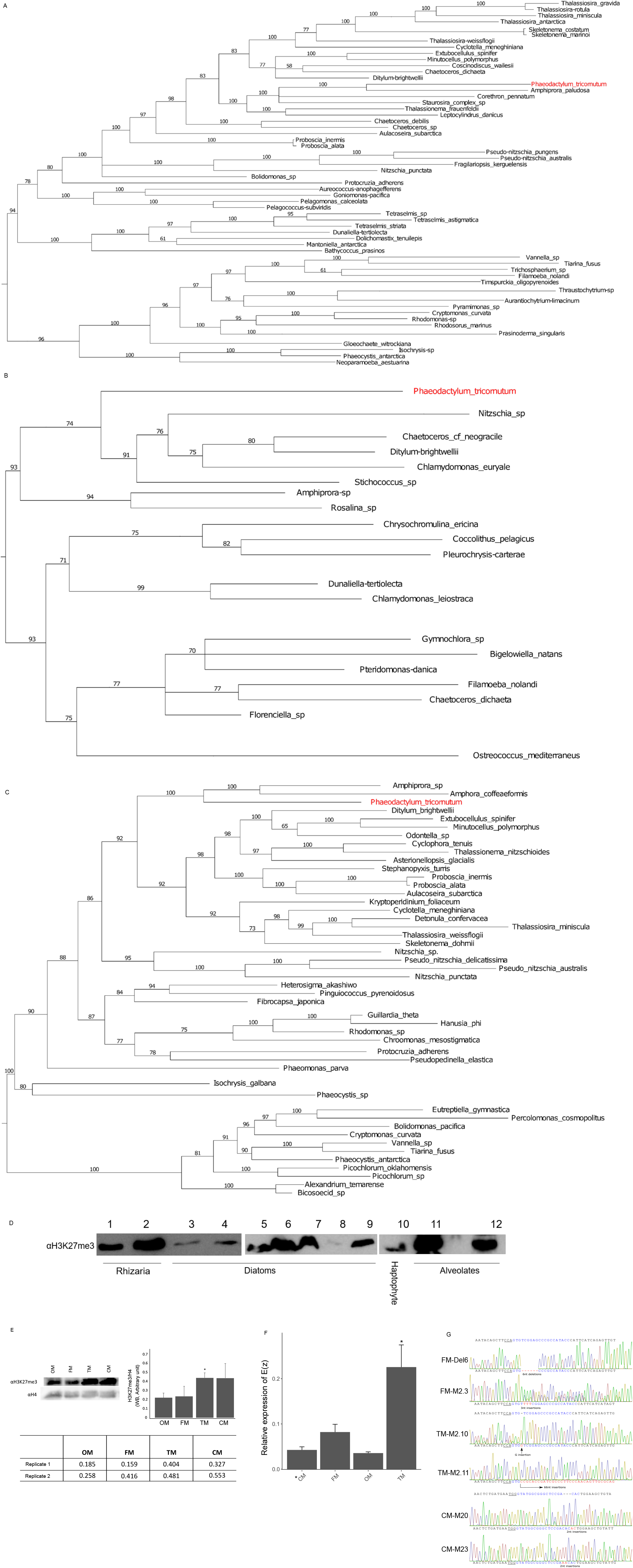
Phylogeny of polycomb complex subunit **A:Enhance of Zest, B:Suz(12,)C:Esc)**. **(D)** Western blots using a monoclonal antibody against H3K27me3 on protein or chromatin extracts of different species representative of the super SAR lineage (1: *Bigelowiella_natans*, 2: *Gymnophora dimorpha*, 3: *Skeletonema marinoi*, 4: *Thalassiosira pseudonana*, 5: *Raphoneis sp*, 6: *Synedra sp*, 7: *Asterionellopsisglacialis*, 8: *Thalassiosira rotula*, 9: *Phaeodactylum tricornutum*, 10: *Isocrhrysis lutea*, 11: *Amhedinium klebselii*, 12: *Amhedinium carteri*). **(E)** Western blot on chromatin extracts from oval (OM), fusiform (FM), triradiate (TM) and cruciform (CM) with a monoclonal antibody against H3K27me3 and a densitometry quantification, Bar plot at up left was quantification of H3K27me3/H4 using pixel calculation by ImageJ from two independent western blot, Table below shows the calculated number of two western at up right. **(F)** Expression of E(z) in OM,FM and TM by RT-QPCR. (**G)** Sequence chromatograms of PCR product from WT cells and CRISPR cas9 mutants of enhancer of zeste showing the different indels in FM, TM and CM.

**Supplementary Figure 2.**
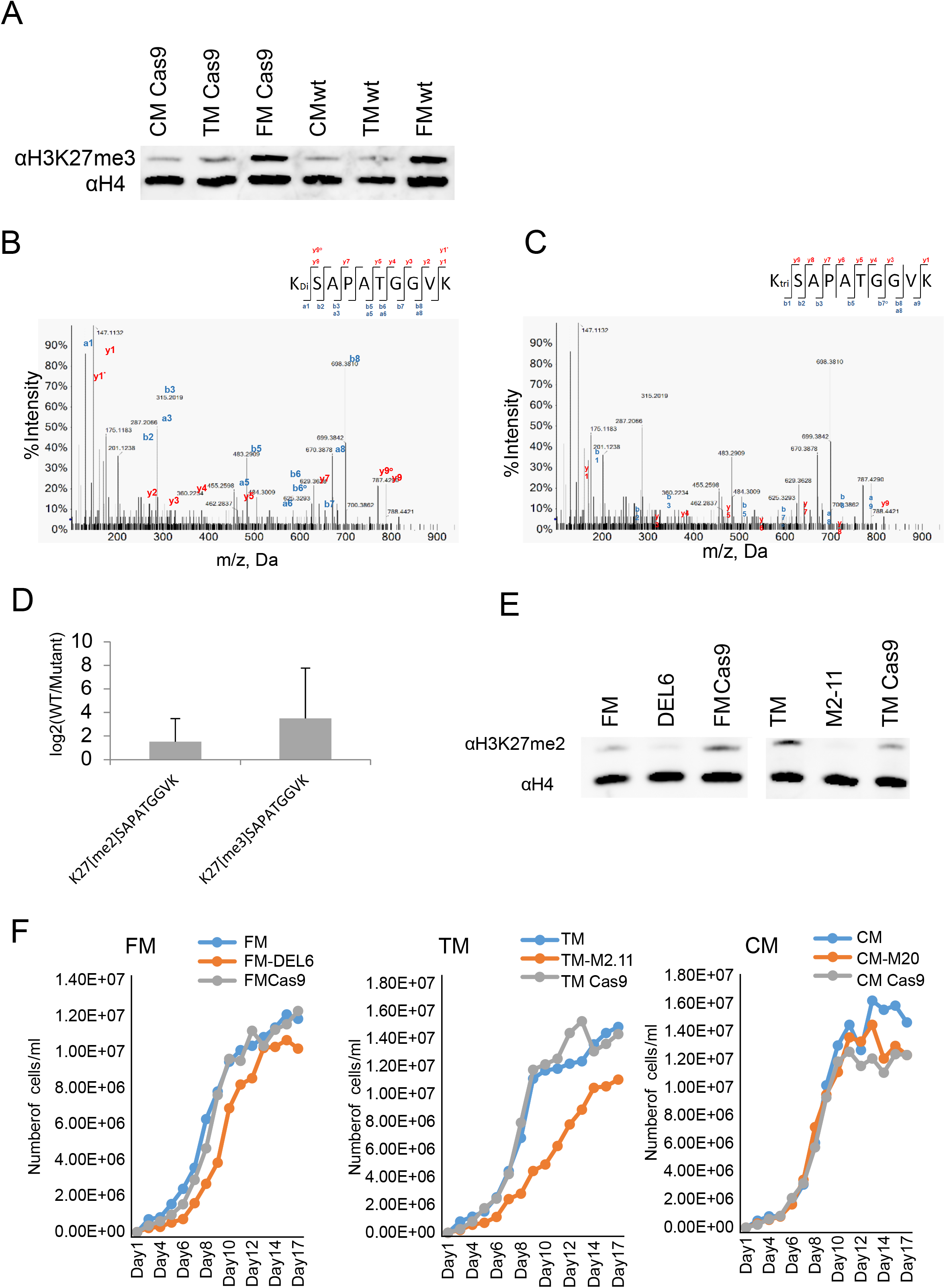
**(A)** Western blot of chromatin extracts of Cas9 control lines from each of FM, TM and CM. H4 histone antibody was used as a loading control. Cas9 control are the empty vector controls containing Cas9 and Shble antibiotic resistance gene. **(B, C)** Mass spectrometry quantification of di and tri-methylation of lysine 27 of histone H3 in both wild type and enhancer of zeste knockout mutant. MS/MS spectrum of the [M + 2H]^2+^ precursor ion of histone H3 (27–36 residue peptide) tri-methylated or di-methylated on K27. Broken bonds above and below sequence denote b and y ions, respectively, that were annotated from the spectrum. **(D)** Abundance of H3 K27 di- and tri-methylated KSAPATGGVK peptide. Y axis shows normalized log2 (WT/Mutant) of the di-methylated and tri-methylated peptides. All measurements have been performed in triplicate, and error bars indicated the 97.5% confidence interval (see supplementary Table 1). **(E)** Western blot of chromatin extracts from wild type FM and TM as well as E(z) knockouts in both backgrounds with a monoclonal antibody against H3K27me2. H4 histone antibody was used as loading control. **(F)** Growth curves of wild type, enhancer of zeste mutants and cas9 control line in each of FM, TM and CM.

**Supplementary Figure 3.**
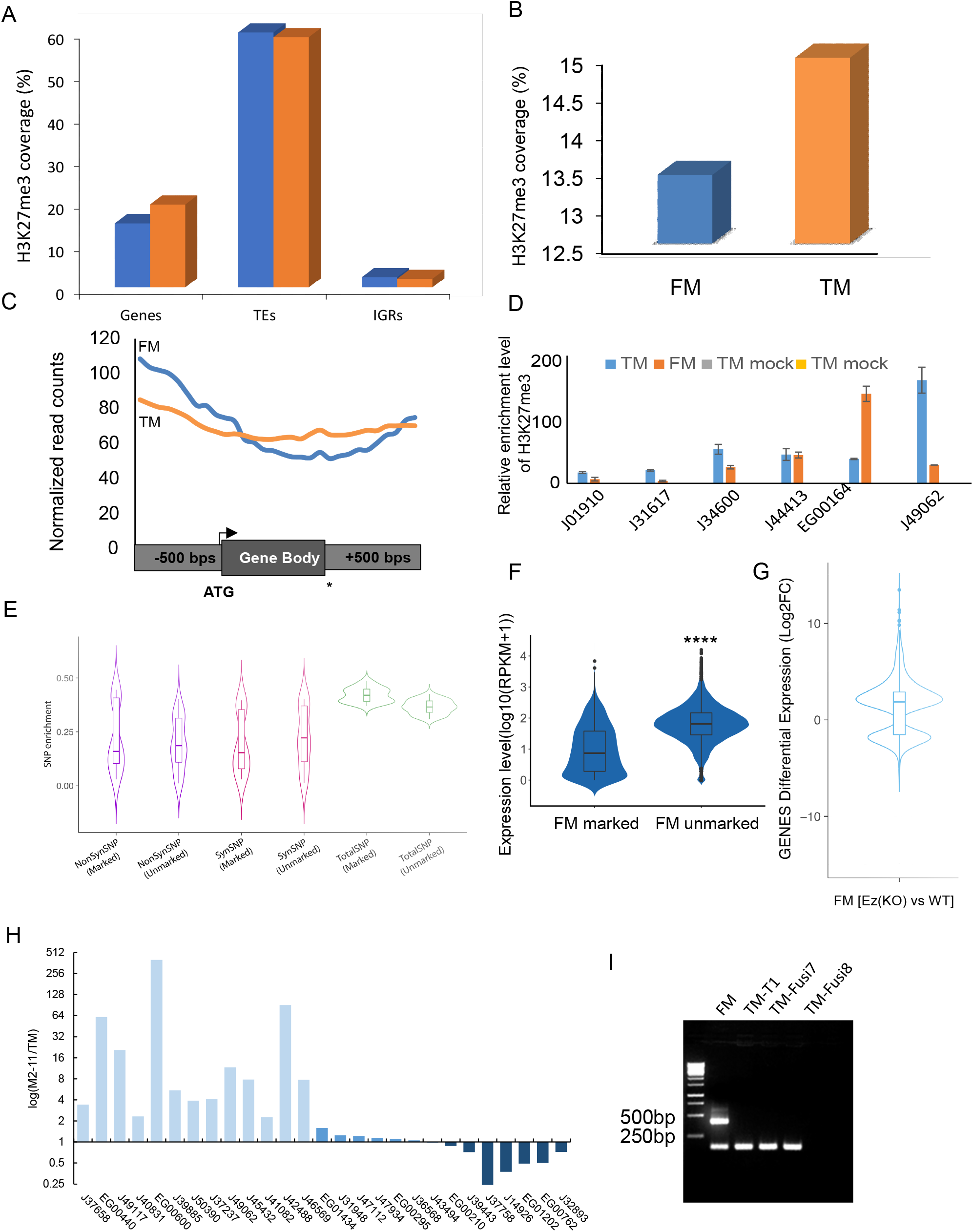
**(A)** Bar plot present the coverage of H3K27me3 on TEs, Genes and Inter generic regions (IGRs) respectively. **(B)** Total genome coverage of H3K27me3 within TM (orange) and FM (blue) showing a higher mapping of H3K27me3 in TM compared to FM. **(C)** Mean distribution of H3K27me3 over 500 bp upstream, genebody, and 500 bp downstream region of all the gene targets in TM (orange line) and FM (blue line). **(D)** Unmarked genes (J01910, J31617), commonly marked genes (J34600, J44413) were chosen as internal controls. FM (EG00164) and TM specifically marked (J49062) genes were used as controls for the reproducibility of independent ChIP-QPCR results. **(E)** violin plot represents SNPs comparison of genes marked by H3K27me3 and unmarked genes. **(F)** violin plot shows the expression level of marked genes and unmarked genes in FM, significant of differences were estimated by two-tailed t-test with P value <0.0001, as denoted by “****”. **(G)** violin plot represents the mean fold change of gene expression in Ez(KO) lines compared to the wild-type (WT). Expression fold change with P value < 0.05 are considered for plotting. **(H)** Relative expression level of H3K27me3 targets genes in the TM and enhancer of zeste knock out M2-11. In the plot, fold change log2(PtM2-11/Pt8Tc) values are shown. **(I)** Gel picture of a molecular marker distinguishing FM and TM and amplifying an insertion in one allele present in FM but absent in TM.

**Supplementary Figure 4.**
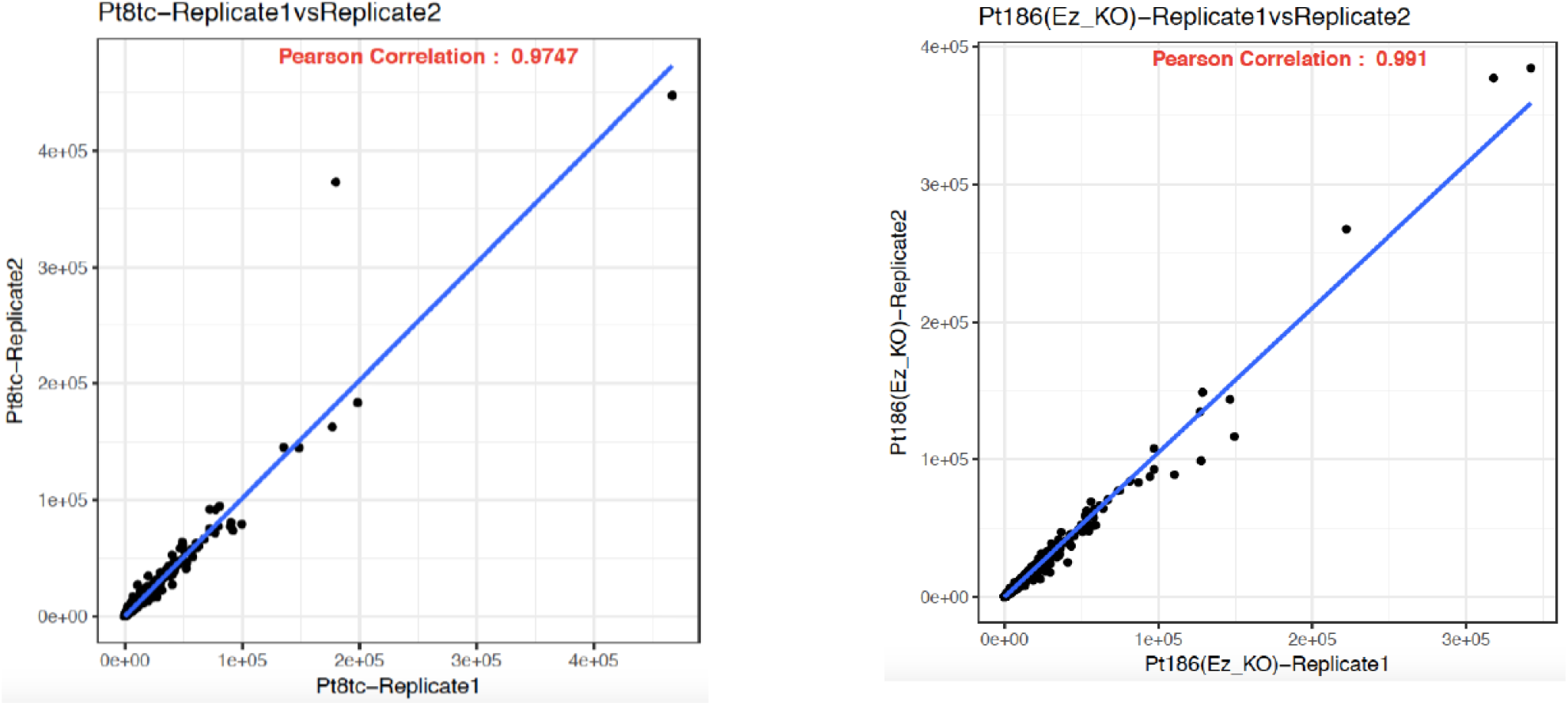
Scatter plots with Pearson correlation coefficient displaying the relationship between TM and E(z) knock out RNA Seq replicates.

**Supplementary Figure 5.**
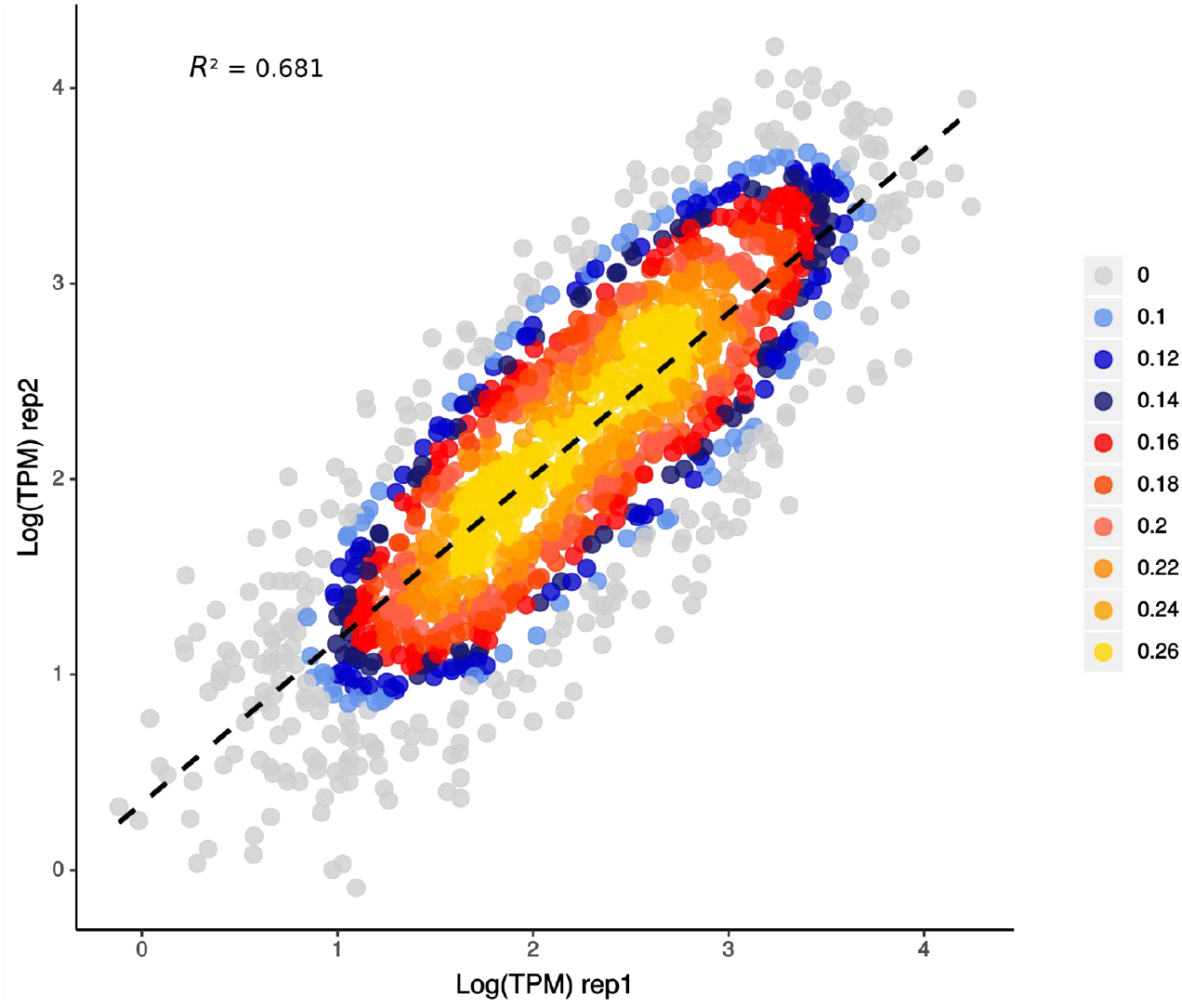
Scatter plots with Pearson correlation coefficient displaying the relationship between TM ChIP-Seq replicates

## Acknowledgements

Hanhua Hu from the Chinese Academy of Science is acknowledged for the gift of CM morphotype.). LT acknowledges funds from the CNRS, the region of Pays de la Loire (ConnecTalent EPIALG project) and Epicycle ANR project (ANR-19-CE20-0028-02). CB acknowledges funding from the ERC Advanced Award ‘Diatomite, the French Government ‘Investissements d’Avenir’ programmes MEMO LIFE (ANR-10-LABX-54), PSL* Research University (ANR-1253 11-IDEX-0001-02). XZ was supported by a PhD fellowship from the Chinese Scholarship Council (CSC-201604910722). Dr. Gu Jie is acknowledged for his help with data analysis. AR was supported by an International PhD fellowship from the MEMO LIFE Program.

## Author contributions

LT and XZ conceived and designed the study. XZ, LT and AFDC performed experiments, AR, LT, XZ, VR, FRJV and OA analysed the data. CC maintained the cultures, XL helped with ChIP seq, BL and DL performed the Mass spec analysis, AG and AV contributed to the bioinformatics analysis, DB and CB contributed to the discussion of the results and text editing. All authors analysed and interpreted the results. LT and XZ wrote the manuscript with input from all the authors.

